# Nucleotide Analogues as Inhibitors of Viral Polymerases

**DOI:** 10.1101/2020.01.30.927574

**Authors:** Jingyue Ju, Shiv Kumar, Xiaoxu Li, Steffen Jockusch, James J. Russo

## Abstract

Coronaviruses such as the newly discovered virus from Wuhan, China, 2019-nCoV, and the viruses that cause SARS and MERS, have resulted in regional and global public health emergencies. Based on our molecular insight that the hepatitis C virus and the coronavirus use a similar viral genome replication mechanism, we reasoned that the FDA-approved drug EPCLUSA (Sofosbuvir/Velpatasvir) for the treatment of hepatitis C will also inhibit the above coronaviruses, including 2019-nCoV. To develop broad spectrum anti-viral agents, we further describe a novel strategy to design and synthesize viral polymerase inhibitors, by combining the ProTide Prodrug approach used in the development of Sofosbuvir with the use of 3’-blocking groups that we have previously built into nucleotide analogues that function as polymerase terminators.

---

The recent appearance of clusters of a new coronaviral infection in Wuhan, China has made headlines, as more and more people have become infected in China and in a number of other countries. Already more than 100 deaths have been ascribed to this virus, and there are fears of additional outbreaks. The virus has been isolated from the lower respiratory tracts of patients with pneumonia, sequenced and visualized by electron microscopy (Zhu et al 2020). The virus, which has received the placeholder name 2019-nCoV, turns out to be a new member of the subgenus sarbecovirus, in the Orthocoronavirinae subfamily, but is distinct from MERS CoV and SARS CoV (Zhu et al 2020). The coronaviruses are positive-sense, single strand RNA viruses, and thus share properties with other single-stranded RNA viruses such as hepatitis C virus, West Nile virus, Marburg virus, HIV virus, Ebola virus, dengue virus, and rhinoviruses.

The coronavirus life cycle is depicted in Fig. 1, which is taken from a review by Zumla et al (2016). Briefly, the virus enters the cell by endocytosis, is uncoated and ORF1a and ORF1b of the + strand RNA is translated to produce nonstructural proteins, including a cysteine protease, a serine protease, helicase and RNA-dependent RNA polymerase (RdRp). The proteases cleave the precursor forms of the helicase and RdRp to form mature, functional molecules. A replication-transcription complex is then formed, which is responsible for making more copies of the RNA genome via a negative-sense intermediate, as well as the structural and other proteins encoded by the viral genome. The viral RNA is packaged into viral coats, attaches to the cell surface, after which exocytosis results in release of viral particles for subsequent infectious cycles. Potential inhibitors have been designed to attack nearly every stage of this process, as indicated in Fig. 1 (Zumla et al 2016). Despite decades of research, no effective drug is currently available to treat such serious coronavirus infections as SARS, MERS, and the symptoms caused by 2019-nCoV.

**Fig. 1:**
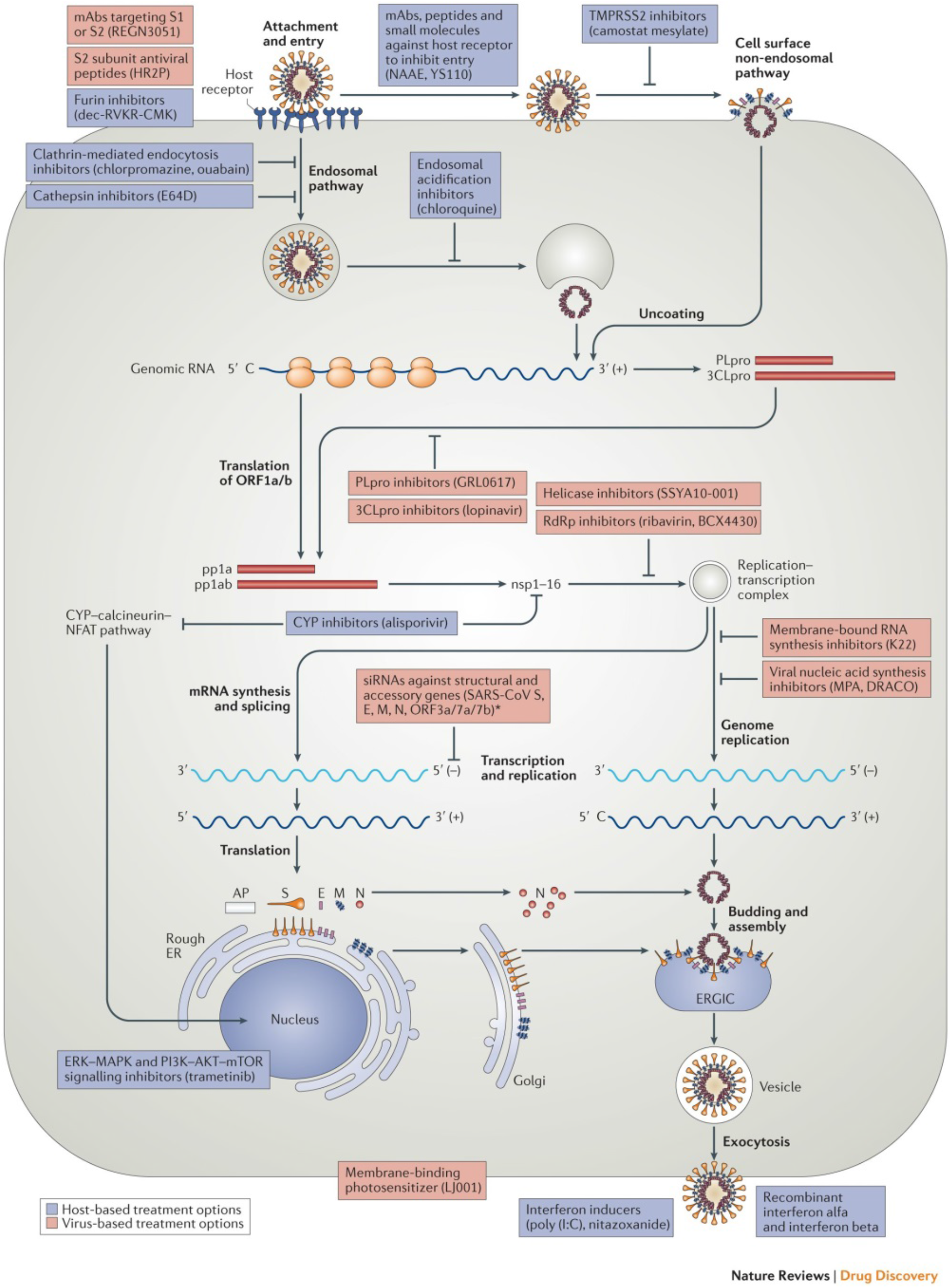
Virus-based and host-based treatment options targeting the coronavirus replication cycle. *From Zumla et al (2016) Nat Rev | Drug Discovery 15:327-347.*

One of the most important druggable targets is the RdRp. Example drugs include Gilead’s sofosbuvir (which is paired with velpatasvir as the FDA-approved drug EPCLUSA), to inhibit the RNA-dependent RNA polymerase of the hepatitis C virus. Sofosbuvir, a pyrimidine nucleotide analogue (Fig. 2) with a blocked phosphate group enabling it to enter infected eukaryotic cells, is a prodrug, which is converted into its active triphosphate form by cellular enzymes (Fig. 3). The activated drug binds in the active site of the RdRp, where it is incorporated into RNA, and due to modifications at the 2’ position, inhibits further RNA chain extension and halts RNA replication. It acts as an RNA polymerase inhibitor by competing with natural ribonucleotides.

**Fig. 2:**
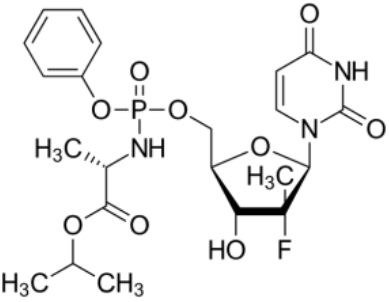
Structure of Sofosbuvir, a component of the FDA-approved drug EPCLUSA.

**Fig. 3:**
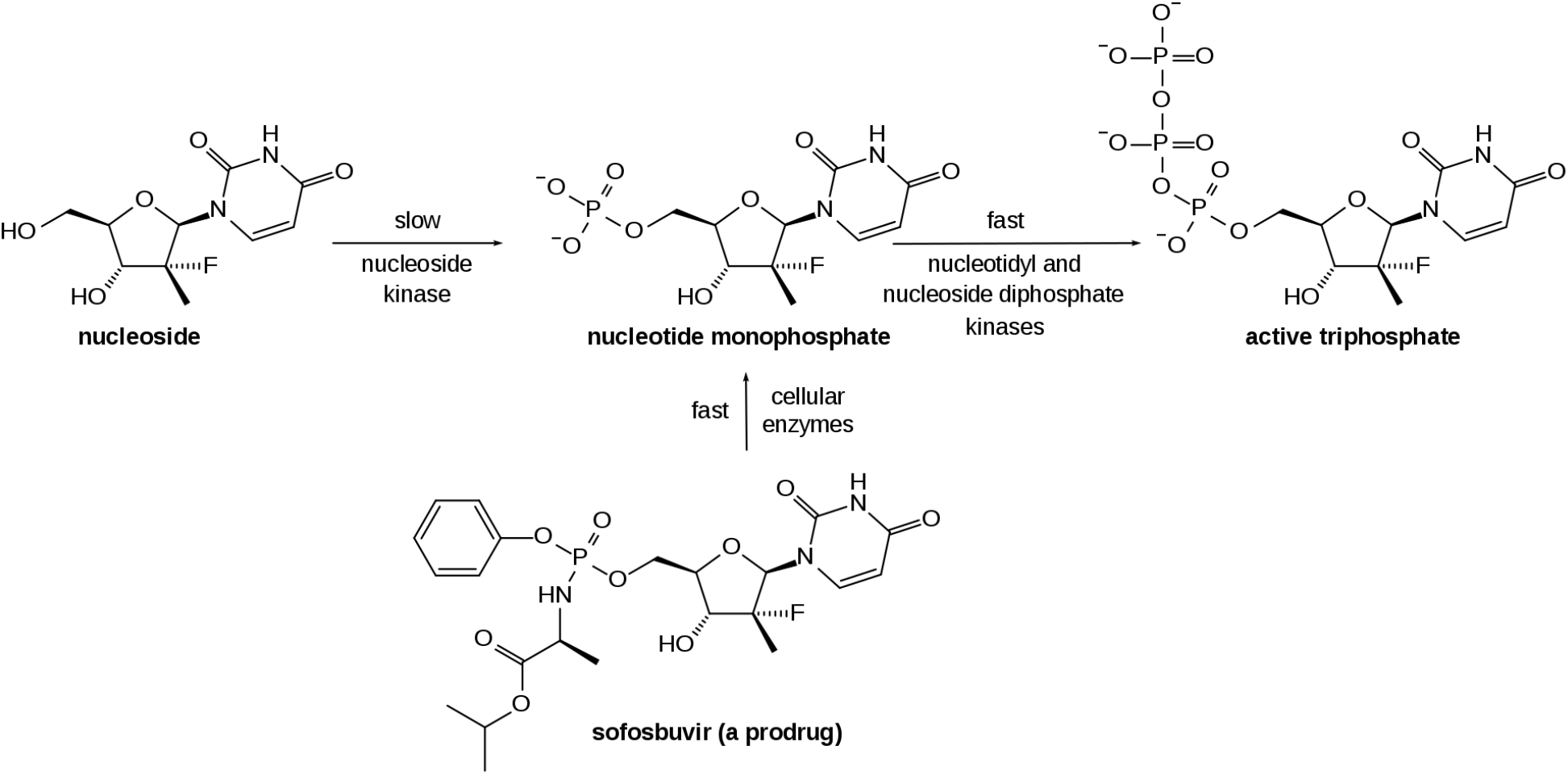
Conversion of sofosbuvir to active drug *in vivo. From https://en.wikipedia.org/wiki/Sofosbuvir.*

Based on our insight that the hepatitis C virus and the coronavirus use a similar viral genome replication mechanism, we reasoned that the FDA-approved drug EPCLUSA for the treatment of hepatitis C will also inhibit coronaviruses, including 2019-nCoV and those responsible for SARS and MERS.

There are many other RNA polymerase inhibitor drugs which are used as antivirals. A related purine nucleotide prodrug, remdesivir (Fig. 4), was developed by Gilead to treat Ebola virus infections, though not very successfully, and is currently being considered for repositioning to treat the 2019-nCoV outbreak (https://www.fiercebiotech.com/biotech/gilead-mulls-repositioning-failed-ebola-drug-china-virus). In contrast to sofosbuvir, both the 2’- and 3’-OH groups are unmodified, but a cyano group at the 1’ position presumably serves to inhibit the RdRp in the active triphosphate form. Additional information on the ProTide Prodrug technology is described by Alanazi et al (2019).

**Fig. 4:**
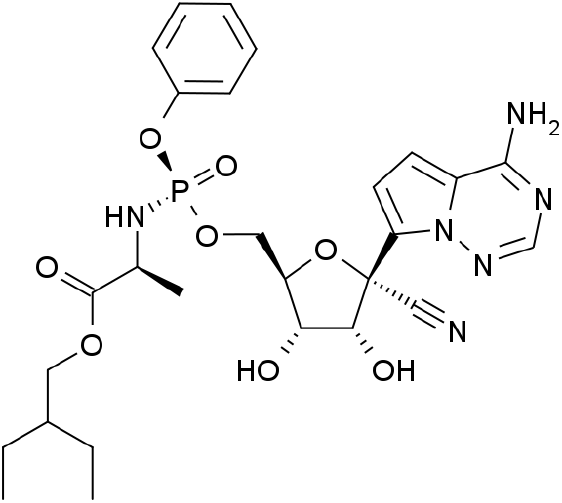
Structure of remdesivir.

A related prodrug analogue developed by BioCryst Pharmaceuticals, BCX4430, also known as galidesivir (Fig. 5), has been shown to inhibit RNA polymerases from a broad spectrum of RNA viruses, notably including the filoviruses (e.g., Ebola, Marburg) in rodents and Marburg virus in macaques (Warren et al 2014). Upon entry into infected cells, BCX4430 is rapidly phosphorylated, and the resulting nucleoside triphosphate serves as an RNA chain terminator.

**Fig. 5:**
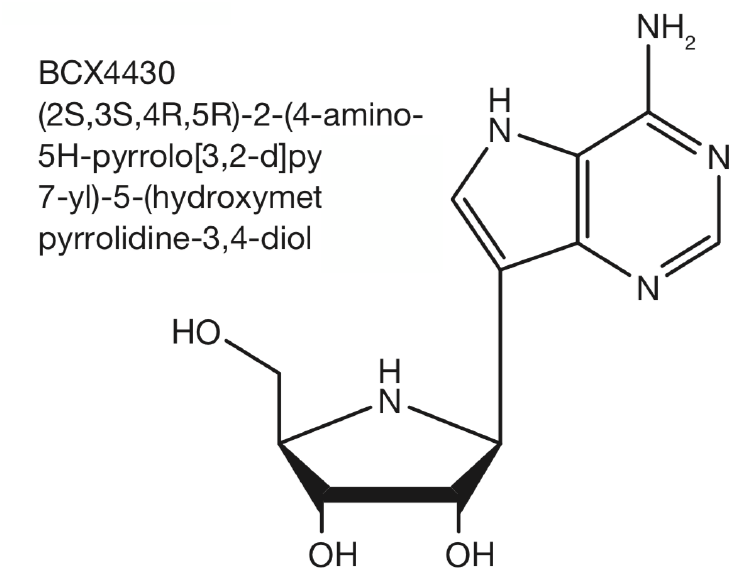
Structure of BCX4430.

Based on the above background information, we describe here a novel strategy to design and synthesize viral polymerase inhibitors, by combining the ProTide Prodrug approach used in the development of Sofosbuvir with the use of 3’-blocking groups that we have built into nucleotide analogues that function as reversible terminators for DNA sequencing (Ju et al 2003, Ju et al 2006, Guo et al 2008). We reasoned that (1) the phosphate masking groups will allow entry of the compounds into infected cells, (2) the 3’-blocking group on the 3’-OH with either free 2’-OH or modifications at the 2’ position will encourage incorporation of the activated triphosphate analogue by viral polymerases but not host cell polymerases, thus reducing any side effects, and (3) once incorporated, further extension will be prevented by virtue of the 3’-blocking group, thereby inhibiting viral replication. These modified nucleotide analogues should be potent polymerase inhibitors and thus active against various viral diseases, including but not limited to the coronaviruses such as 2019-nCoV, and the strains causing SARS and MERS. Once incorporated, our newly designed nucleotide analogues containing 3’ blocking groups will permanently block further viral genome replication. This is in contrast to other nucleotide analogue-based viral inhibitors that have a free 3’ OH group, which have the possibility of allowing further polymerase extension, enabled by viral mutations. For the same reason, at high concentrations, nucleotide analogue-based viral inhibitors with free 3’ OH groups have the potential of being incorporated by host polymerases.

All RNA viruses are known to mutate at a high frequency, due to the low fidelity of the viral polymerase, resulting in the development of resistance to treatment. The promiscuous nature of the viral polymerase will allow incorporation of our newly designed nucleotide analogues as anti-viral agents.

Examples of nucleotide analogues designed by us to satisfy these criteria are provided in Fig. 6, and strategies for their synthesis in Figs. 7–9; Fig. 10 shows the activation of these prodrugs to form triphosphate analogues (same as for sofosbuvir in Fig. 3) that should be incorporated and inhibit the coronavirus and other RNA virus polymerases.

**Fig. 6:**
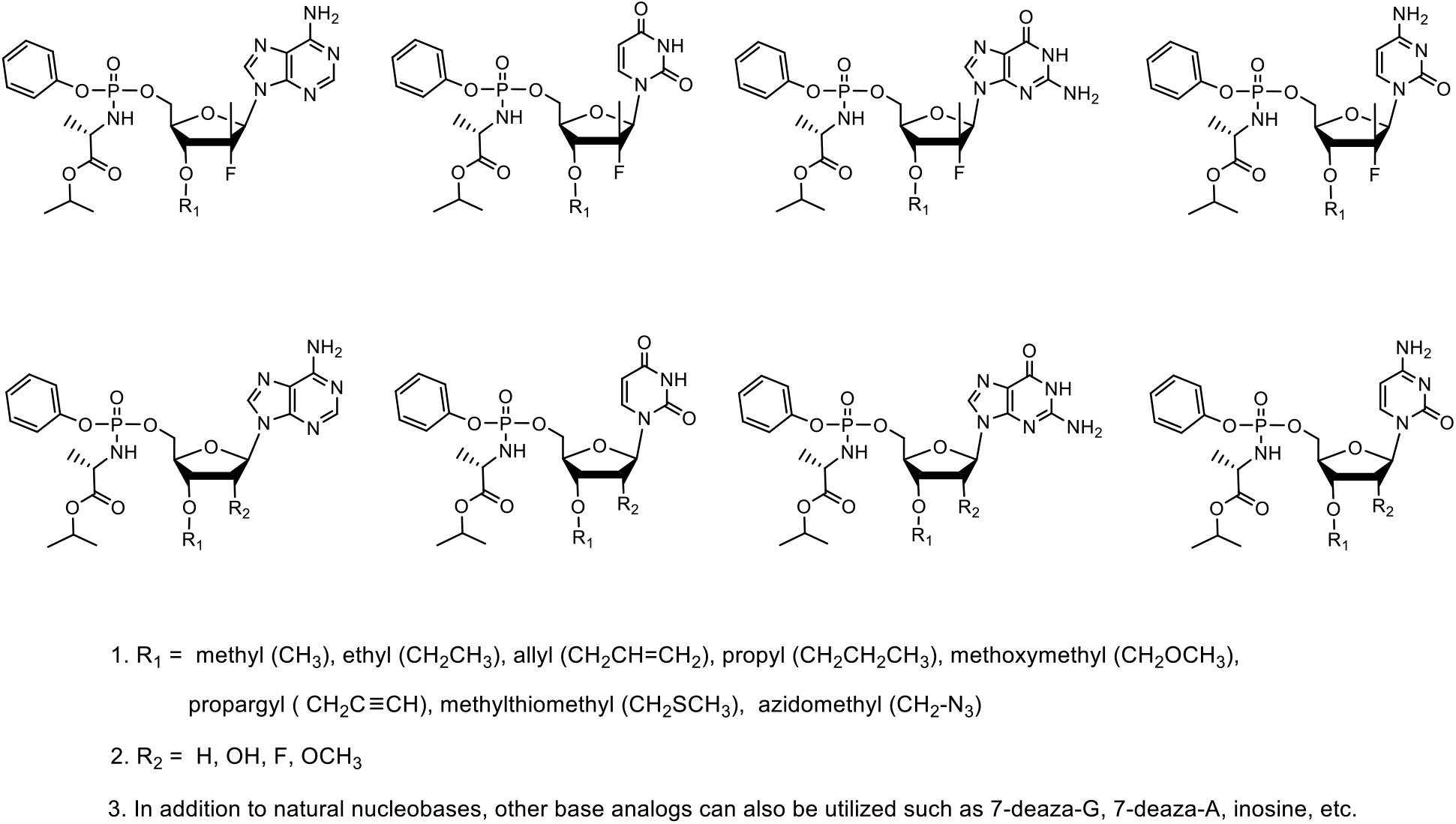
Example structures of nucleotide analogues as viral polymerase inhibitors with masked phosphate group, unmodified or modified at 2’ position, and blocking groups on the 3’ position.

**Fig. 7:**
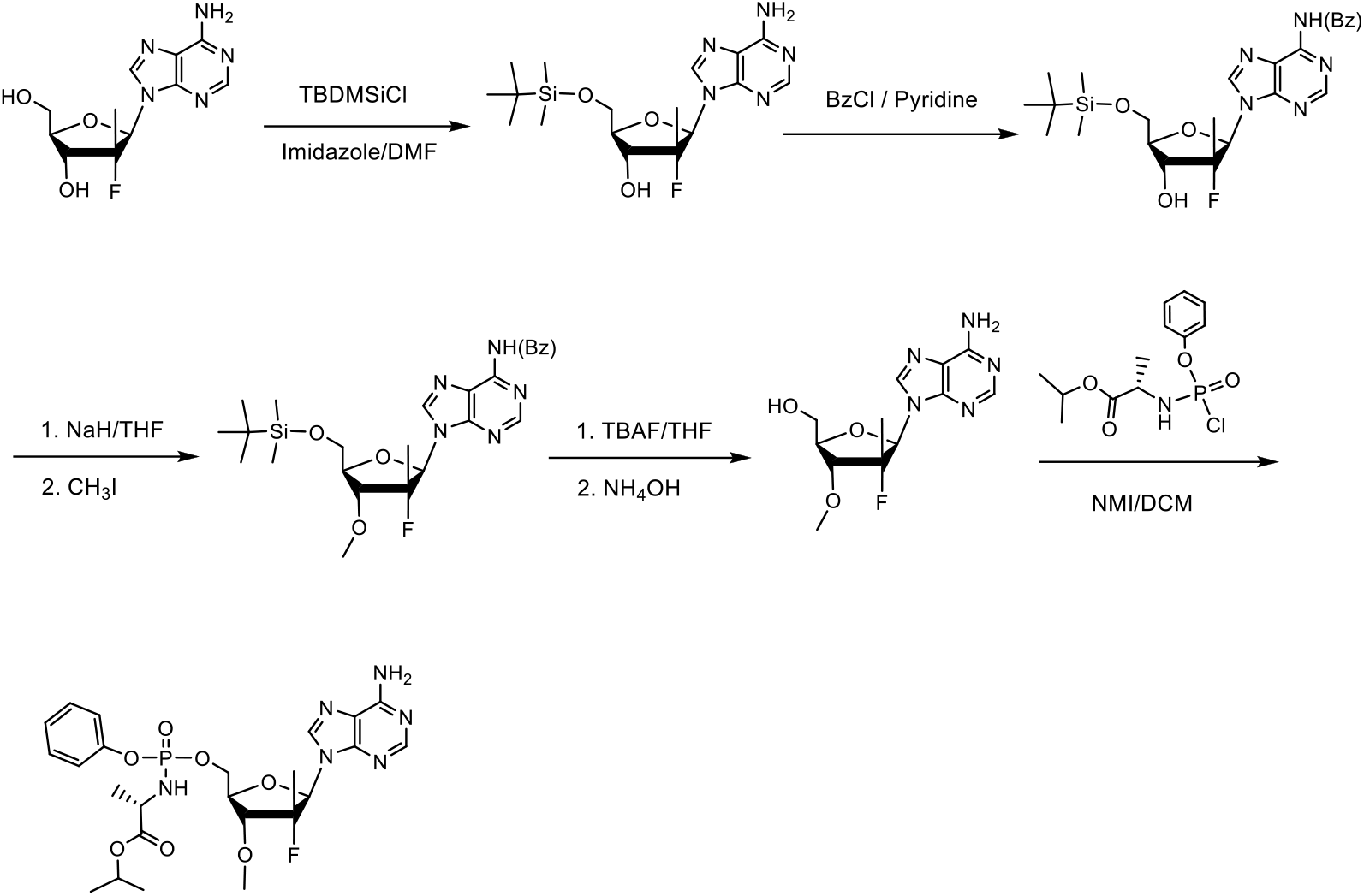
Synthesis of 3’-O blocked nucleoside phosphoramidate analogues: β-D-2’-deoxy-2’-α-fluoro-2’-β-C-methyl-3’-O-methyladenosine nucleoside phosphoramidate as example.

**Fig. 8:**
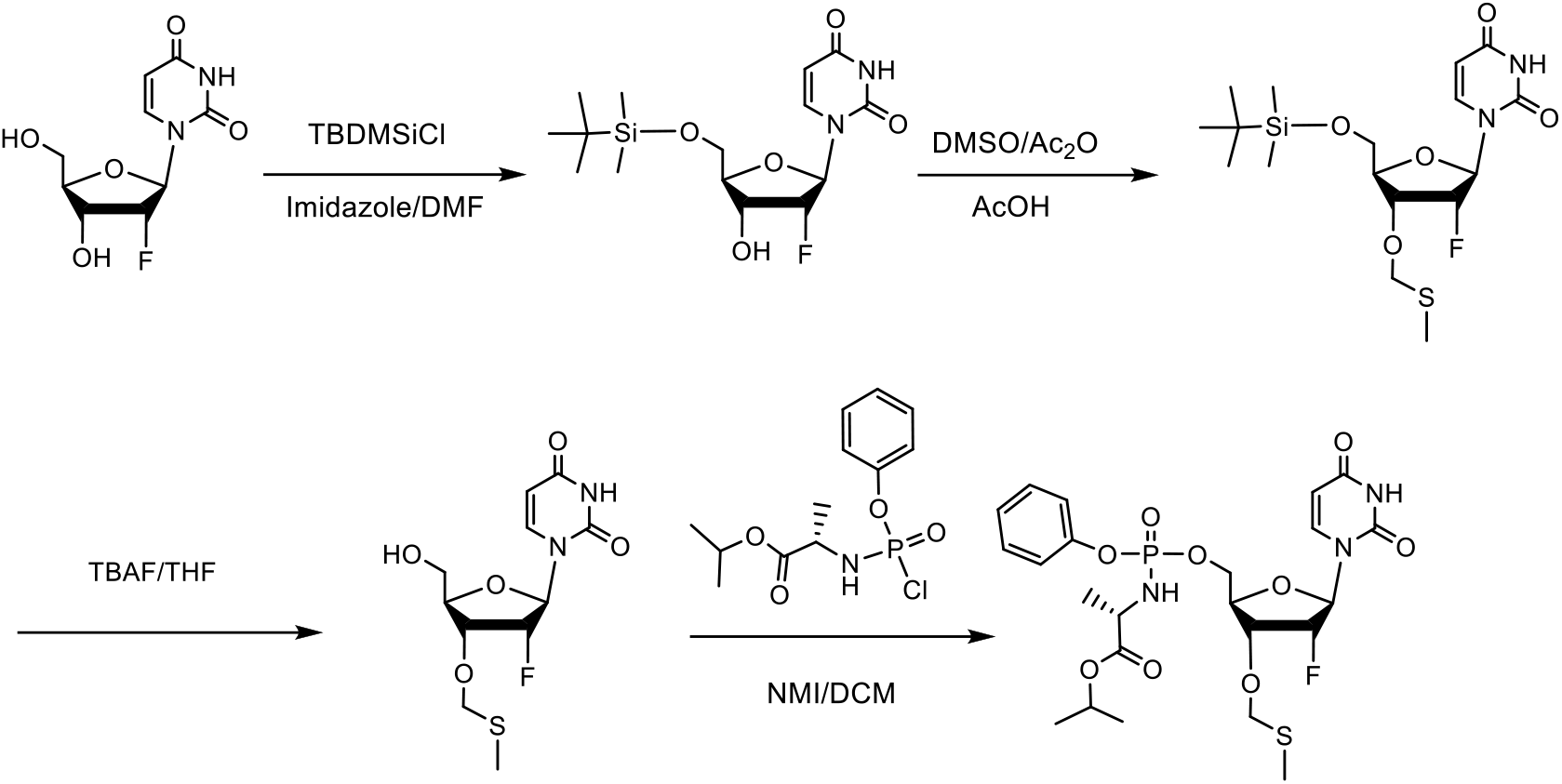
Synthesis of 3’-O blocked nucleoside phosphoramidate analogues: β-D-2’-deoxy-2’-α-fluoro-3’-*0*-methylthiomethyluridine nucleoside phosphoramidate as example.

**Fig. 9:**
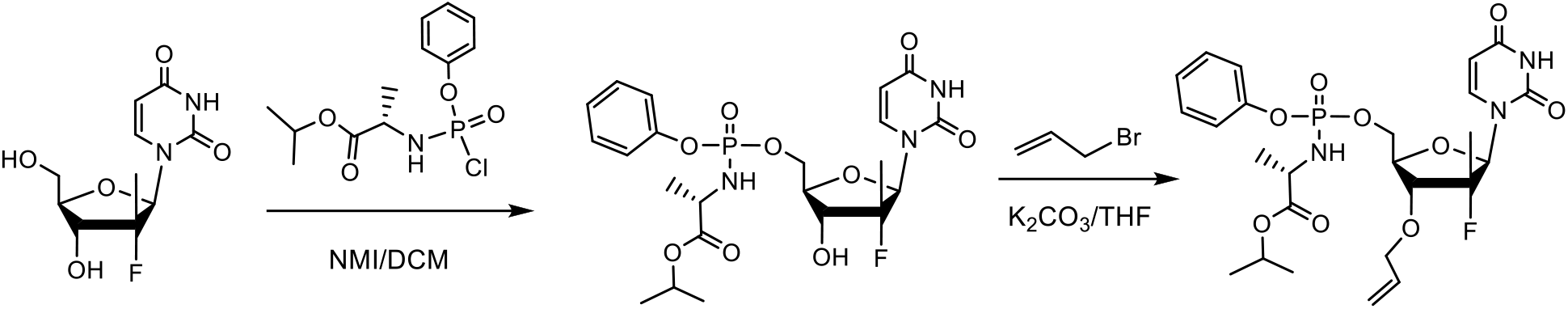
Example synthesis of β-D-2’-deoxy-2’-α-fluoro-2’-β-C-methyl-3’-*O*-allyluridine nucleoside phosphoramidate.

**Fig. 10:**
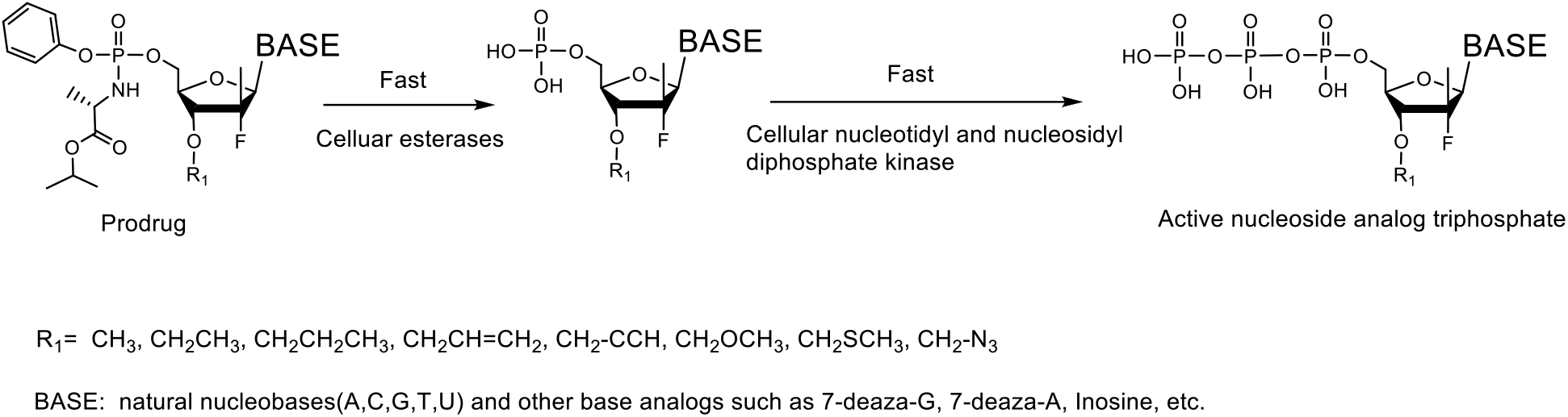
Mode of activation of 3’-blocked nucleotide prodrug precursors illustrated in Fig. 6 by cellular enzymes.

Synthesis of 3’-*O*-blocked nucleoside phosphoramidate prodrugs can be carried out starting from 2’-modified nucleosides (Ross et al 2011). First, both the 5’-OH and the exocyclic amino group of the base will be protected. Then the 3’-OH will be derivatized with a variety of blocking groups, including methyl, ethyl, propyl, allyl, propargyl, methoxymethyl, methylthiomethyl, azidomethyl, etc., such as those listed in Fig. 6, following established methods (Ju et al 2006, Guo et al 2008). After deprotection, the free 5’-OH is derivatized to afford the corresponding phosphoramidates by treatment with freshly prepared chlorophosphoramidate reagent in the presence of N-methyl imidazole (Sofia et al 2010). Fig. 7 and Fig. 8 show example synthetic routes for the 3’-methoxy and 3’-O-methylthiomethyl nucleoside phosphoramidate analogs, respectively. Alternatively, starting from a 2’-modified nucleoside, the 5’-OH can be derivatized first to give 5’-phosphoramidate nucleotides, followed by 3’-OH derivatization to afford 3’-O blocked nucleoside phosphoramidate analogues. Fig. 9 shows an example synthetic route for 3’-allyl nucleoside phosphoramidate analogues.

## ACKNOWLEDGMENTS

This research is supported by Columbia University, which has filed a patent application on the work described in this manuscript.

